# In-vivo targeted tagging of RNA isolates cell specific transcriptional responses to environmental stimuli and identifies liver-to-adipose RNA transfer

**DOI:** 10.1101/670398

**Authors:** J. Darr, M. Lassi, Archana Tomar, R. Gerlini, F. Scheid, M. Hrabě de Angelis, M. Witting, R. Teperino

## Abstract

Bio-fluids contain various circulating cell-free RNA transcripts (ccfRNAs). The composition of these ccfRNAs varies between bio-fluids and constitute tantalizing biomarker candidates for several pathologies. ccfRNAs have also been demonstrated as mediators of cellular communication, yet little is known about their function in physiological and developmental settings and most works are limited to in-vitro studies. Here, we have developed iTAG-RNA, a novel method for the unbiased tagging of RNA transcripts in mice in-vivo. We used this method to isolate hepatocytes and kidney proximal epithelial cells-specific transcriptional response to a dietary challenge without interfering with the tissue architecture, and to identify multiple hepatocyte-secreted ccfRNAs in plasma. We also identified transfer of these hepatic derived ccfRNAs to adipose tissue, where they likely serve as a buffering mechanism to maintain cholesterol and lipid homeostasis. Our findings directly demonstrate in-vivo transfer of RNAs between tissues and highlight its implications for endocrine signaling and homeostasis.

## Introduction

Little is known about the biological function of circulating cell-free RNAs (**ccfRNA**). Found to be associated with exosomes, lipoproteins, ribonucleoproteins and more, these transcripts can be isolated and sequenced from multiple bio-fluids such as plasma, lymph, cerebral fluids, breast milk and more [1, 2]. ccfRNAs are directly implicated in the development of several pathologies including cancer and obesity [3-5] and are intensively studied as disease biomarkers [6, 7]. Despite this, the role they play in physiological and developmental settings and in mediating cell-to-cell communication remains largely unknown. In-vitro, a growing number of works demonstrate the relevance of RNA based cellular communication [8-11], however in-vivo evidence is still limited. This discrepancy is partly due to the difficulties posed to tracking ccfRNAs from transcriptional source to potential sites of action in-vivo. Indeed the tools available to study ccfRNAs in physiological settings are limited and very few studies attempt to tackle this problem directly. One work found evidence to suggest that the majority of circulating miRNAs originate in adipose tissue and that some of the adipose derived miRNAs may play a role in the regulation of liver Fgf21 levels [12]. However, this work focuses on miRNA and does not directly demonstrate transfer of RNAs between tissues nor directly identify adipose secreted RNAs.

Transfer of miRNAs was also demonstrated between epithelial cells of the caput epididymis to maturing spermatozoa, leading to a shift in sperm RNA content during its maturation [13]. This study made use of 4-thiouracil-tagging (TU-tagging) [14] combined with SLAM-Seq [15] to demonstrate loading of miRNAs transcribed in caput epididymis into maturing spermatozoa. TU-tagging entails cell-type specific expression of uracil phosphoribosyltransferase (UPRT) and administration of 4-thiouracil, with the assumption that only cells expressing UPRT would incorporate 4-thiouracil into transcribing RNA. Thio-RNA can then be purified and used for downstream gene expression analyses, or alternatively combined with SLAM-Seq to identify labeled transcripts. TU-tagging has proven useful in several additional systems [14, 16, 17], however, given endogenous [18] and alternative [19] pathways for uracil incorporation, the labeling specificity in this method remains unclear. In addition, as is demonstrated in Herzog et. al [15] and by Sharma et. al [13], labeling with TU-tagging of PolI and PolIII transcripts is inefficient, rendering tRNAs and ribosomal transcripts unlabeled.

Indeed there are only a limited number of techniques enabling in-vivo targeted labeling of RNAs. In addition to TU-tagging, 5-ethynylcytosine-tagging (EC-tagging) [20] is a new method, which utilizes cell-type specific co-expression of cytosine deaminase (CD) with UPRT to achieve RNA labeling with 5-ethynyluridine (5EU) following administration of 5-ethynylcytosine. Both TU and EC tagging use cre-recombination to express the relevant enzymes in a tissue specific manner and stochastic expression from the cre-promoter may lead to unwanted expression of the enzymes in different tissues [21]. Finally, one recently developed method called Mime-seq allows for cell type specific labeling of microRNA [22]. In this method, tissue specific expression of a plant derived methyltransferase mediates a 3′-terminal 2′-O-methylation of microRNAs that, when combined with a methylation dependent library construction, allows for sequencing of tissue specific microRNAs. Mime-seq allows labeling of miRNAs alone leaving other RNA biotypes unlabeled. Given the need for a technique that allows for a Cre-independent and unbiased labeling of total RNA transcription in-vivo, we developed iTAG-RNA **[For In-vivo Targeted Tagging of RNA]**. This method incorporates mouse genetics with a novel uridine analog and an established RNA labeling chemistry to allow tagging of total RNA in target cells in-vivo. Using iTAG-RNA we are able to identify transcriptional re-programming of hepatocytes in-vivo following an acute high fat diet stress and to enrich for and identify hepatocyte derived plasma ccfRNAs. Moreover, we are able to identify RNA-based liver-to-adipose RNA transfer. These liver derived ccfRNAs include variable coding and non-coding RNAs such as miRNAs and tRNAs. Among the miRNAs transferred from liver to adipose tissue we find mir-33, mir-10b and mir130a, which target major regulators of cholesterol and lipid efflux and bio-synthesis such as Srebf1 [23], Abca1 [23, 24], Ppara [25] and Pparg [26] respectively.

Our study demonstrates for the first time an unbiased technique that allows labelling, tracking and quantification of variable types of ccfRNAs from their transcriptional source to downstream tissues, in which they can potentially act to regulate expression of target genes. We demonstrate RNA-based liver-to-adipose transfer of a myriad of RNA transcripts and their response to an environmental challenge. The continued identification and characterization of RNA based signaling in-vivo is imperative for the understanding of developmental, physiological and pathological processes, and can aid in the future development of relevant disease biomarkers.

## Results

### Small molecule design and genetic approach for targeted in-vivo labeling of RNA

5-Ethynyl Uridine **(5EU)** is a synthetic uridine analogue extensively used in RNA turnover studies [27-29]. The nitrogenous base contains an alkyne group that can be covalently linked to an azide group using a simple copper mediated reaction called click chemistry [30, 31]. This synthetic base has been demonstrated to incorporate into transcribing RNA in place of uridine and have little to no biological effects thereafter [27]. Following administration to mice, 5EU is readily taken up by cells with no regard to cell identity, depending to some extent on the administration method and dosage used [27]. Here, we present a novel method for the targeted in-vivo delivery of 5EU. To achieve this, we designed a ‘pro-drug’ of the 5EU base **(HD5EU)** that is based on the ‘Hep-Direct’ pro-drug design [32, 33] (Figure 1a). This design was developed to target small molecules and nucleotide analogues to the human CYP3A4 enzyme and several small molecules of this design have been or are currently in clinical studies [34-36]. The human CYP3A4 enzyme catalyzes an oxidative cleavage of the HD5EU small molecule which, following a spontaneous beta-elimination, results in the formation of 5EU mono-phosphate that can then be incorporated into transcribing RNA (Figure 1a). HD5EU was synthesized by Chiroblock GmbH, the identity of the final product was validated using MS, p-NMR and h-NMR and the molecule’s purity was assessed at over 98% (Sup. Figure 1a-d).

**Figure 1.**
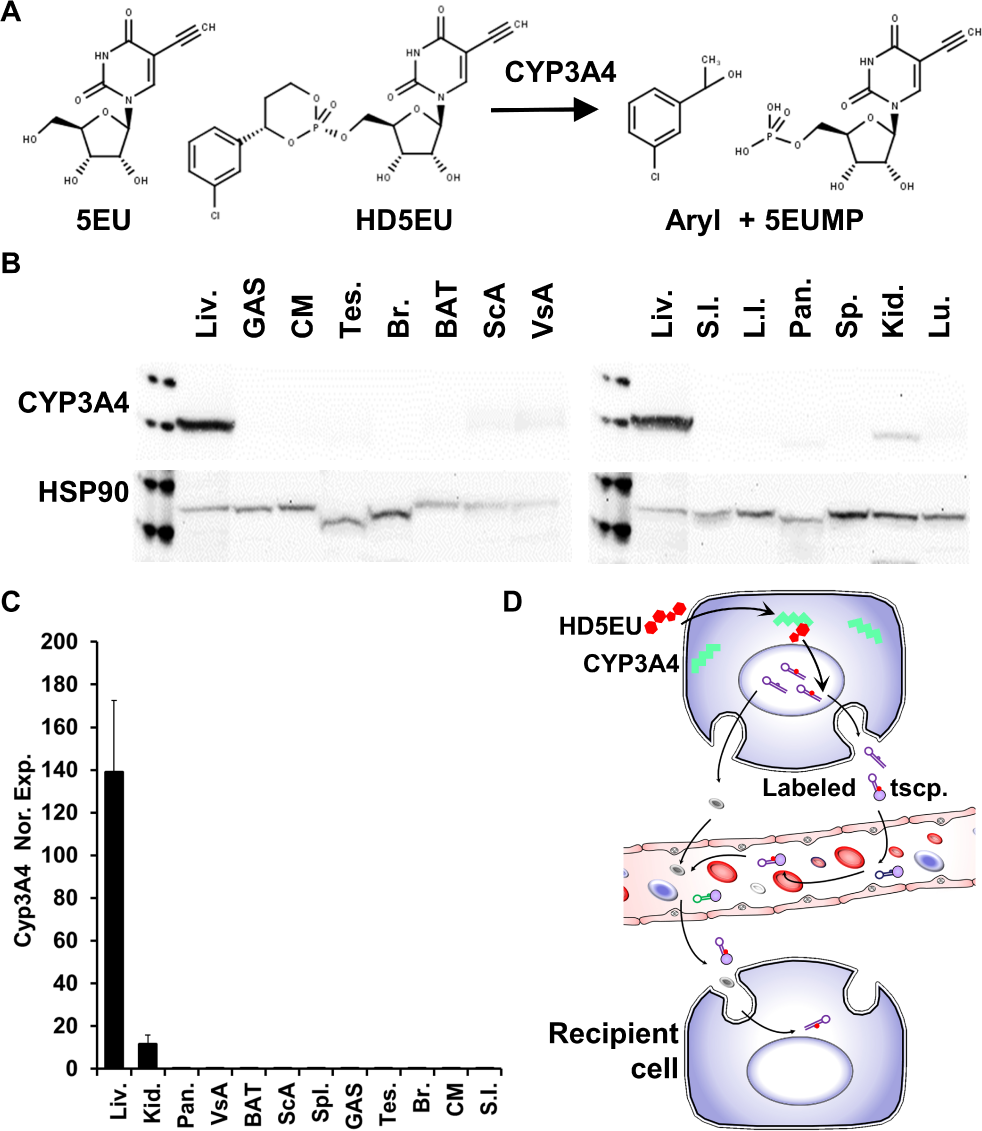
HD5EU small molecule design and CYP3A4 expression pattern. **A)** HD5EU small molecule and metabolite structure relative to 5EU. **B)** W.B. depicting tissue expression pattern of CYP3A4 in humanize CYP3A4 cluster deleted mice. HSP90 serves as loading control. GAS = Gastrocnemius muscle; CM = Cardiac Muscle; BAT = Brown adipose tissue; ScA = Subcutaneous white adipose tissue; VsA = Visceral white adipose tissue; S.I. = Small intestine; L.I. = Large intestine. **C)** qRT-PCR validating liver and kidney specific expression of CYP3A4. Error-bars for standard-deviation of 3 biological repeats. **D)** Administration of the HD5EU small molecule to cells expressing CYP3A4 allows metabolism to 5EU and labelling of total RNA. Labeled transcripts are then secreted to the extracellular matrix and can be identified in bio-fluids and recipient cells.

In addition to the HD5EU small molecule we took advantage of the published humanized liver specific CYP3A4 mouse line **FVB/129P2-*Cyp3a13***^**tm1Ahs**^ **Del(*5Cyp3a57-Cyp3a59*)**^**1Ahs**^ **Tg(*APOE*-CYP3A4)**^**A1Ahs**^ obtained from Taconic [37, 38]. These humanized mice **(hCYP3A4)** express the human CYP3A4 enzyme under a modified Apolipoprotein E (APOE) promoter and are stably knocked out for nine homologous murine genes, thus leaving the human enzyme as the sole member of the enzyme family to be expressed in a Cre-independent, tissue-specific manner in-vivo. In keeping with published data on the activity of the modified ApoE promoter [39], qRT-PCR and WB analyses demonstrate restricted expression of the human Cyp3a4 enzyme to liver and kidney (Figure 1b-c). As such, upon administration of the HD5EU small molecule to the humanized CYP3A4 mice, we expect the molecule to be metabolized to bioavailable 5EU mono-phosphate exclusively in cells expressing CYP3A4, namely hepatocytes and kidney proximal renal epithelial cells, thus allowing in-vivo targeted labelling of transcription and identification of secreted transcripts in bio-fluids upon pull-down of 5EU labelled RNA (Figure 1d).

### CYP3A4 is necessary for in-vitro and in-vivo metabolism of the HD5EU small molecule

To test and validate the metabolism of the HD5EU small molecule, we first isolated primary hepatocytes from hCYP3A4 mice. Following an 8 hour treatment of primary hepatocytes with 1mM HD5EU or 5EU, nuclear staining similar to 5EU labeling is clearly evident following click-it fluorescent staining (Figure 2a). When pre-treated with Azamulin, a highly selective CYP3A4 inhibitor [40], HD5EU treated primary hepatocytes no longer demonstrate nuclear staining, in contrast to 5EU treated primary hepatocytes, where nuclear staining is unaffected by Azamulin pretreatment (Figure 2a). These results indicate that while 5EU is still readily incorporated into transcribing RNA in the nucleus, nuclear staining in HD5EU treated cells is dependent upon CYP3A4 activity.

**Figure 2.**
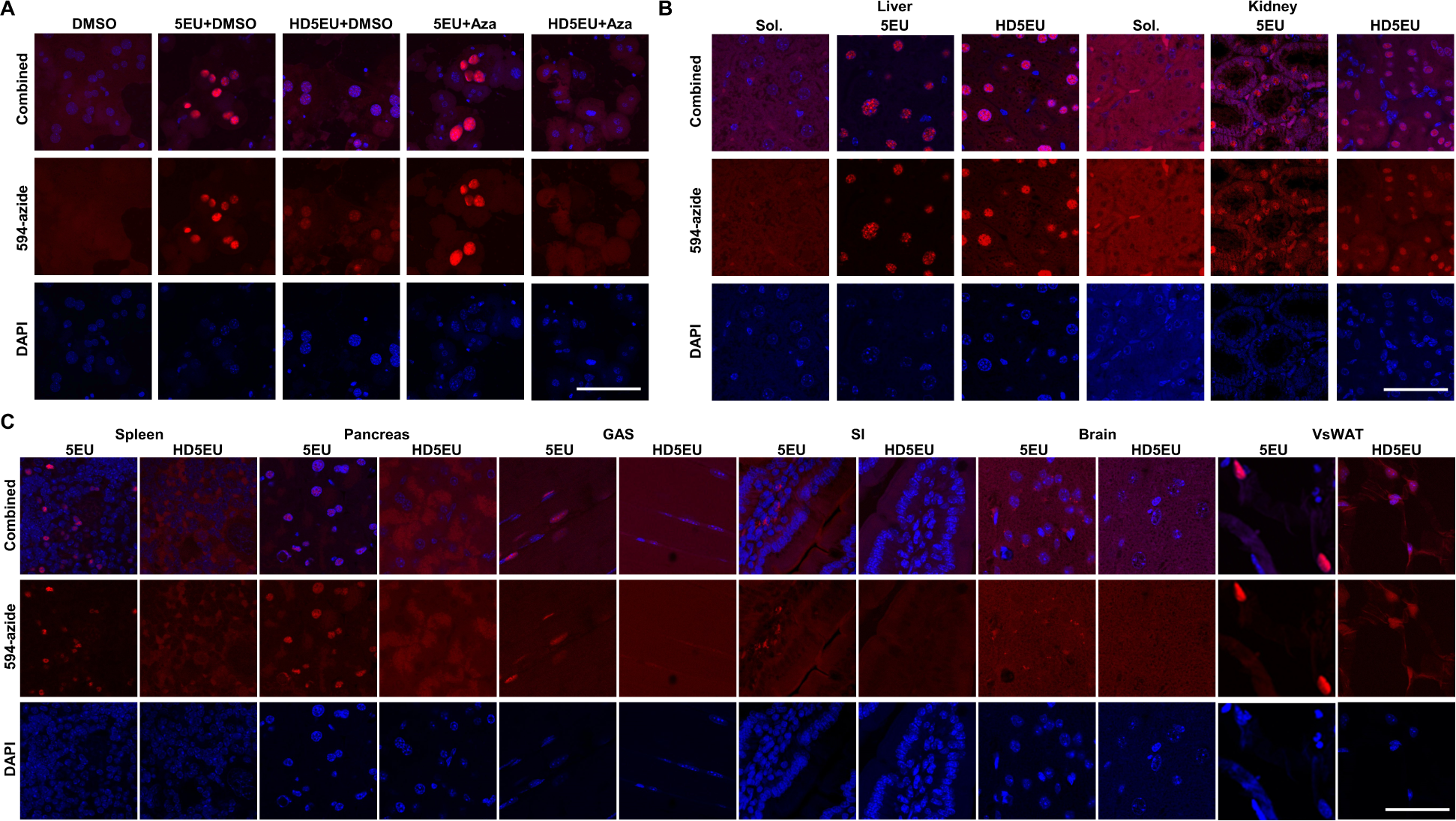
Tissue specific staining evident with HD5EU is dependent on CYP3A4 activity. **A)** Primary hepatocytes demonstrate fluorescent nuclear staining of nascent RNA transcription following 8 hours of 5EU and HD5EU treatment. Azamulin treatment hinders fluorescent nuclear staining in HD5EU treated cells but not 5EU treated cells. 594-azide used for click-it staining. **B)** Following 2 hours of HD5EU administration to mice, in-vivo fluorescent nuclear staining is restricted to hepatocytes and epithelial cells in the kidney and absent from other tissue as evident in panel C and supplementary figure 2. Sol. = solvent for HD5EU. **C)** A similar treatment with 5EU in-vivo results in fluorescent nuclear staining of cells in multiple tissue including; Spleen, Pancreas, Muscle, Small Intestine, Brain and VsWAT. Consecutive injections did not change the observed staining pattern as no tissue apart from liver and kidney demonstrated positive staining. Scale Bar = 50µM. For negative controls see supplementary figure 2.

We administered HD5EU to humanized CYP3A4 mice and already 2 hours following administration found robust nuclear staining in hepatocytes and kidney epithelial cells but in no other tissue examined. This in contrast to mice administered with 5EU where nuclear staining was evident in multiple tissues (Figure 2b-c and Sup. Figure 2a). Of note, animals administered with HD5EU did not demonstrate any visible side effects. In addition we could not detect any signs of DNA damage or apoptosis in the liver during the different treatment regimens as demonstrated by staining for phosph-P53 and cleaved Caspase-3, supporting HD5EU as a non-toxic agent (Sup Figure 2b).

### Mass-Spectrometric validation and quantification of 5EU incorporation into RNA following HD5EU treatment

To validate that nuclear staining evident in-vitro and in-vivo following HD5EU treatment is indeed indicative of 5EU incorporation into transcribing RNA, we adopted the mass-spectrometry method described by **Su et. al [41]**. Using a column with a smaller inner diameter and lower flow rates to improve the response of individual nucleotides, we were able to identify a wide range of unmodified and modified nucleotides **(Sup. Table 1)**. 5EU (m/z 269.0768) was well separated from the potential interfering 13C-Adenosine isotope (Adenosine m/z 268.1040, 13C-Adenosine m/z 269.106541) in standard samples (Figure 3a). Due to mass spectrometric settings all nucleotides show a prominent in-source fragment, which corresponds to the neutral loss of the ribose (Figure 3b, denoted as [M-132+H]^+^).

**Figure 3.**
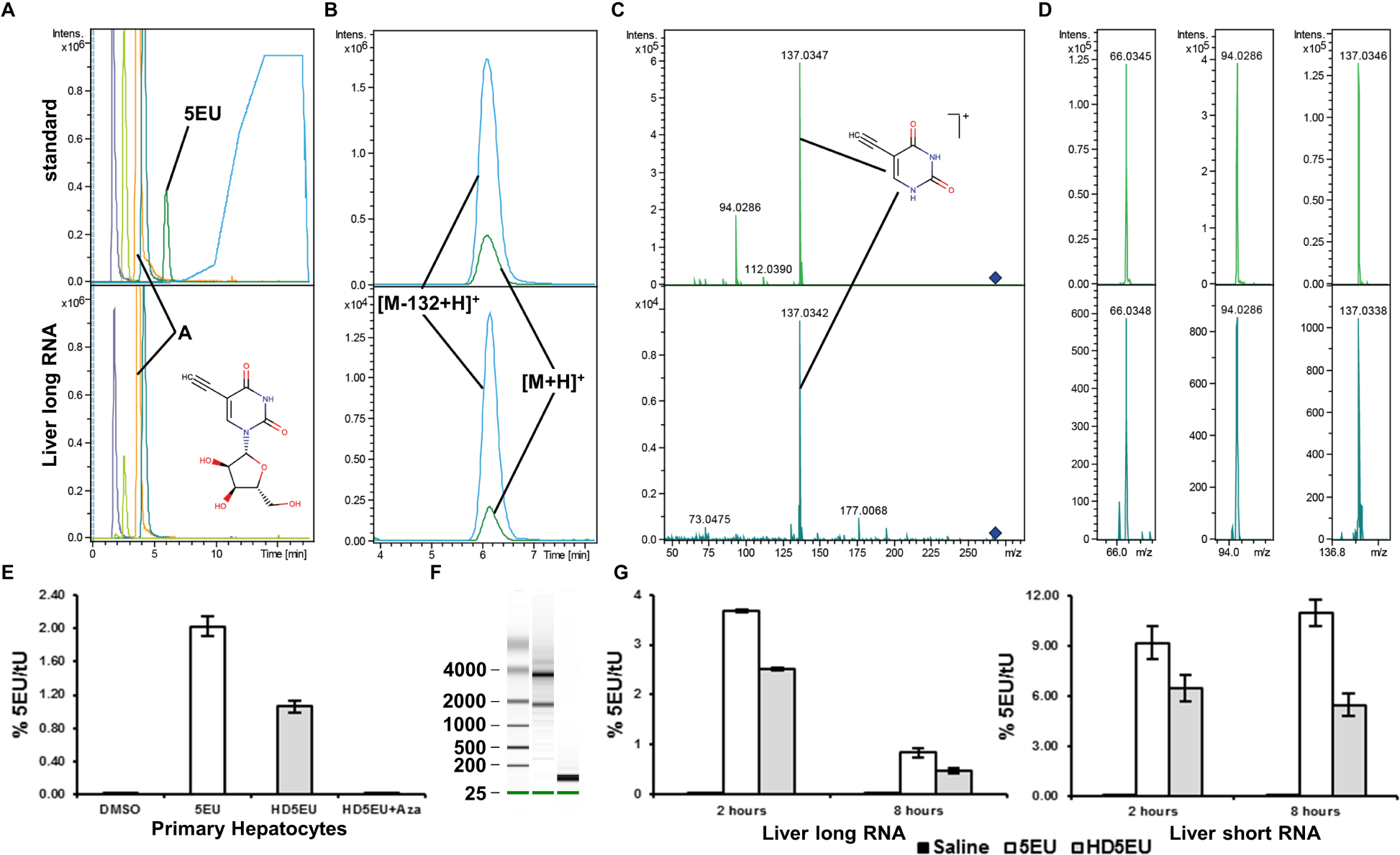
MS validation and quantification of 5EU in liver RNA of HD5EU treated mice. **A)** Extracted ion chromatograms from A, C, G, U and 5EU and gradient slope. **B)** Close up of extracted ion chromatograms of 5EU [M+H]^+^ and 5EU in-source fragment [M-132+H]^+^. **C)** Tandem MS spectra at 20 eV of standard and sample indicating main fragment [M-132+H]^+^. **D)** Further fragments at 40eV used for identification of 5EU in RNA samples. **A-D** upper panels depict standard, lower panels depict liver long RNA from HD5EU treated mice. **E)** Relative quantification of 5EU in primary hepatocytes treated for 8h with the indicated compounds. Azamulin treatment inhibits CYP3A4 mediated HD5EU metabolism to 5EUMP and its subsequent incorporation into RNA. **F)** Bioanalyzer image depicting isolation of liver long and short RNA. **G)** Relative quantification of 5EU in liver derived long / small RNA following indicated time after 5EU / HD5EU administration to mice. Error bars indicated standard deviation calculated for 3 biological replicates.

Tandem mass spectrometry validated that 5EU is present in RNA extracted from the liver of HD5EU treated mice (Figure 3a-d). The main fragment at 20eV collision energy was the neutral loss of ribose ([M-132+H]^+^) consistent with the observed in-source fragment (Figure 3b-c). Further fragments of the remaining nucleoside fragment were observed under higher collision energy of 40 eV (Figure 3d). To quantify 5EU incorporation into RNA we used the prominent in-source fragment of 5EU due to the low abundance of 5EU in biological samples, as this in-source fragment was up to 3-5-fold higher than the intact molecule. In-vitro, Azamulin treatment of primary hepatocytes inhibited HD5EU metabolism and incorporation into transcribing RNA (Figure 3e), in-line with observed fluorescent staining (Figure 2a). In-vivo, we could detect and quantify 5EU incorporation into both short (less than 200bp) and long RNA isolated from the liver of HD5EU treated mice, 2 hours following the administration of the compound (Figure 3f-g). 8h following HD5EU administration 5EU was still detectable in long RNA (though it could not be accurately quantified as it was below quantification limit), whilst only a moderate reduction was detected in short RNA.

Taken together, these results confirm that HD5EU is metabolized in a CYP3A4 dependent manner to 5EU, which is then incorporated into transcribing RNA.

### Robustness and reproducibility of RNA precipitation

2 hours following administration of HD5EU, 5EU-containing liver and kidney transcripts can be biotinylated and pulled-down for next generation sequencing. We persistently failed to generate any amplified libraries following pull-down of unlabeled RNA isolated from liver, plasma or kidney of saline treated control mice (Sup. Figure 3a-e). In addition, following a 2 hour treatment with HD5EU we failed to generate libraries following biotinylation and pull-down from plasma and from additional tissues of the HD5EU-treated mice, this in contrast to liver and kidney where expected library amplicons were generated (Sup. Figure 3a-e). This result was consistent for both poly-A enriched and small RNA libraries and suggests biotinylation to be specific for 5EU containing transcripts. Technical replicates of pull-down libraries demonstrated a high degree of correlation between themselves, supporting the technical robustness and reproducibility of the method (Sup. Figure 3f, spearman correlation coefficient = 0.95).

To further assess the levels of non-specific RNA pull-down, we prepared a 10:1 mixture of non-labeled small RNAs from *S. Cerevisiae* with labeled small RNAs derived from mouse liver. Library construction and sequencing of this mixture following RNA pull-down demonstrated highly effective depletion of yeast RNA compared to input (Figure 4a). These results demonstrate RNA pull-down to be highly selective to biotinylated RNA and non-specific RNA precipitation to be extremely low to not detectable.

**Figure 4.**
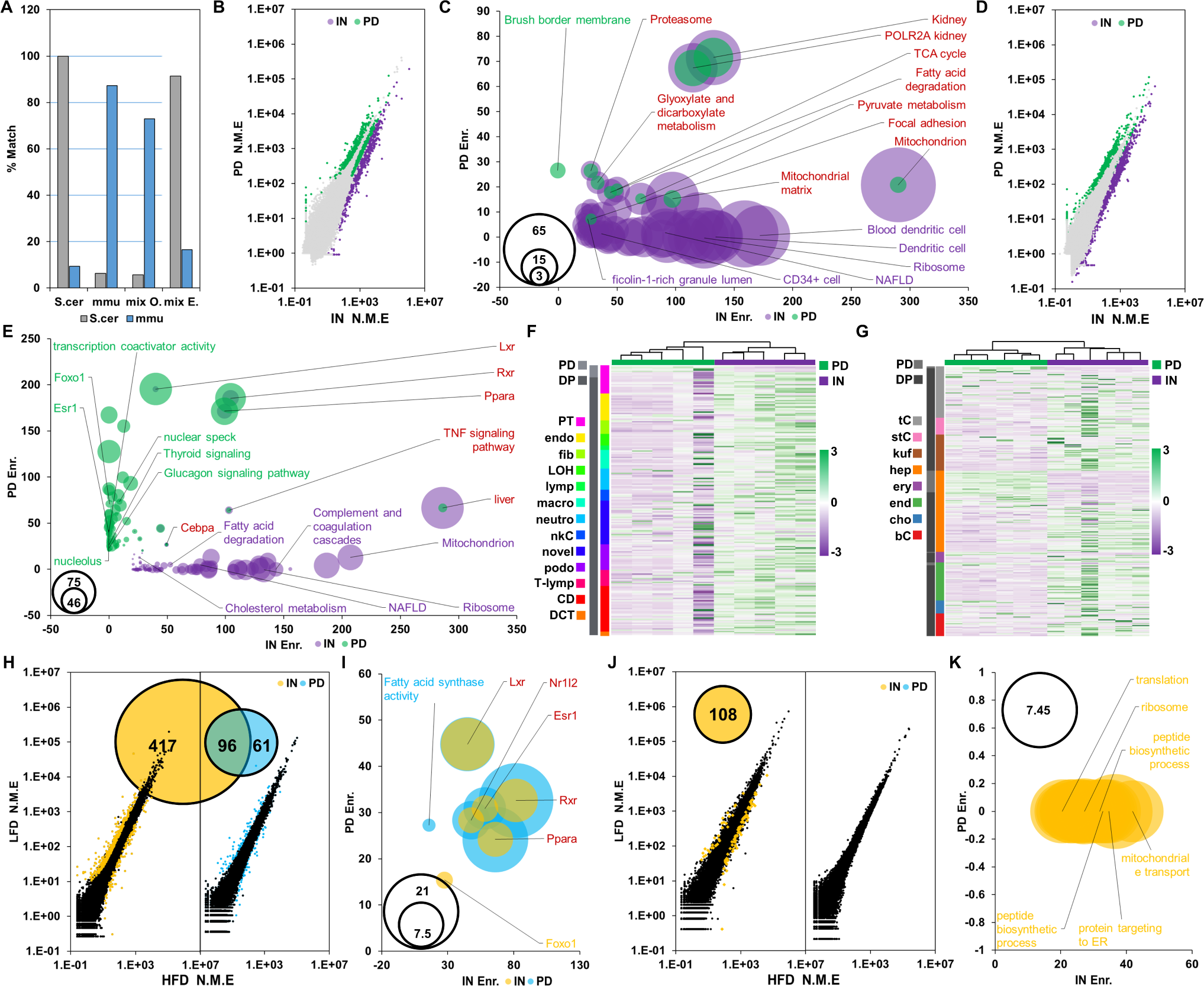
mRNA pull-down enriches for specific cell populations in vivo. **A)** Pull-down of a 10:1 mixture of unlabeled yeast RNA with labeled RNA from mouse liver specifically depletes unlabeled yeast RNA (gray) whilst enriching for the labelled mouse RNA (blue). Compare observed (mix O.) ratios of genomic alignment to expected ratios (mix E.) **B)** Scatter plot visualizing pulled-down (green) and depleted (purple) poly-A RNA from mouse kidney. **C)** GO and gene set enrichment analysis for pulled-down (green) and depleted (purple) protein coding transcripts in kidney. An enrichment for brush boarder membrane proteins, a unique feature in the kidney of proximal epithelial cells, is evident in pulled-down transcripts. Bubble size proportional to –log_10_ of adj. PV. **D)** Scatter plot visualizing labelled pulled-down (green) and depleted (purple) poly-A RNA from mouse Liver. **E)** GO and gene set enrichment analysis for pulled-down (green) and depleted (purple) protein coding transcripts in liver. Bubble size proportional to –log_10_ of adj. PV. **F)** Markers for various cell types found in the kidney and their relative expression in our input and pull-down poly-A RNA libraries. Markers adopted from Park J. et. al. [53]. PT = proximal tubule; endo = endothelial cells; fib = fibroblasts; LOH = Loop of Henle; lymp = lymphocytes; macro = macrophages; neutro = neutrophils; nkC = natural killer cells; novel = novel cell type; podo = podocytes; T-lymp = T-lymphocytes; CD = collecting duct; DCT = distal convoluted tubule. PD = pulled-down, DP = depleted. IN = Input. **G)** Markers for various cell types found in the liver and their relative expression in our input and pull-down poly-A RNA libraries. tC = t-cells; stC = stellate cells; kuf = kuffer cells; hep = hepatocytes; ery = erythrocytes; end = endothelial cells; cho = cholangiocytes; bC = b-cells. Markers adopted from MacParland et. al. [54]. PD = pulled-down, DP = depleted. IN = Input. **H)** Venn diagram demonstrating the overlap between differentially expressed genes in input liver mRNA (orange) and pull-down liver mRNA (Blue) with the corresponding scatterplot. HFD normalized mean expression on the X-axis, LFD normalized mean expression on the Y-axis. **I)** Differentially expressed genes in input kidney mRNA (orange) with the corresponding scatterplot. HFD normalized mean expression on the X-axis, LFD normalized mean expression on the Y-axis. No DEG detected in pull-down mRNA. Bubble size proportional to –log_10_ of adj. PV. **J)** GO and gene set enrichment analysis for dietary induced differentially expressed protein coding genes identified in pull-down (blue) and input (orange) liver libraries. **K)** GO and gene set enrichment analysis for dietary induced differentially expressed protein coding genes identified in input kidney libraries. Bubble size proportional to – log_10_ of adj. PV.

### RNA labelling in-vivo uncovers tissue architecture and stress induced transcriptional reprogramming

We continued to examine if in-vivo labeling enriches for transcriptional programs of specific cellular populations within complex tissues in-vivo, such as the proximal renal epithelial cells in the kidney and hepatocytes in liver, and whether detection of environmentally induced transcriptional reprogramming is possible. To this end we fed mice with high fat (**HFD**) or control low fat (**LFD**) diets for two weeks. Following this acute HFD exposure, which is expected to alter the transcriptional program in the liver [42], we administered HD5EU 2 hours before sacrificing. We then continued to generate poly-A RNA libraries from kidney and liver input and pull-down RNA.

Following mapping with the STAR aligner [43], transcript quantification using HTSeq-count and differential pull-down analyses using the NOISeq package [44], we defined pulled-down transcripts as those whose abundance can be estimated with a high degree of confidence to be at least half of the abundance observed in input (i.e. at least 50% of the gene’s transcripts are labeled with a probability cut-off of 0.975) (Figure 4b-g, **Sup. table 2**).

In kidney, where proximal renal epithelial cells are labelled, GO annotation and gene set enrichment analysis using Enrichr [45] demonstrated an enrichment for genes localized to the brush boarder membrane, along with a few more general terms found enriched also in input such as mitochondria, focal adhesion and genes specific to or highly expressed in the kidney (Figure 4b-c, **Sup. table 3**). The brush boarder membrane in the kidney is a unique feature of proximal renal epithelial cells [46], and among pulled-down transcripts, the Solute Carrier Family 9 member A3 (Slc9a3) is one of its specific markers. Slc9a3 is the sodium–hydrogen antiporter 3, which is highly expressed in the proximal tubule and allows active transport of sodium to the cell. Terms uniquely enriched in genes depleted following pull-down include genes associated with ribosomal function, genes specific to or highly expressed in CD34 positive cells, dendritic cells as well as genes associated to neutrophil ficolin granules. CD34 positive cells are likely endothelial cells found in glomeruli and blood vessels in both human and mice but absent from tubules [47, 48]. In liver, enrichment for identified liver targets of the nuclear receptors PPARA, LXR and RXR is evident in both depleted and pulled-down transcripts. Pulled-down transcripts demonstrate additional enrichment for identified liver targets of transcription factors such as Foxo1, Clock and Nucks1 and for transcripts localizing to nuclear speckles and nucleoli (Figure 4d-e, **Sup. table 4)**. LXR and RXR are implicated in lipid metabolism and heterodimerize to regulate gene expression. Their transcriptional upregulation is associated with increased hepatic lipogenesis [49]. PPARA instead, binds long chain free fatty acids and is a central regulator of lipid metabolism. It heterodimerizes with RXR or LXR to regulate mitochondrial and peroxisomal fatty acid oxidation [50-52].

Published single-cell sequencing from kidney [53] and liver [54] supports depletion of genes associated with irrelevant cell types in both organs. Using published single-cell data, we compared the top ranking genes defined in each study to be cluster specific markers to our pull-down enrichment results. In kidney, we find the majority of cluster markers to be depleted following pull-down of kidney poly-A RNA, apart from a small subset of markers for proximal tubule cells (Figure 4f). In liver, this trend continued and from the list of cluster specific markers, 80% of those identified as enriched following liver poly-A RNA pull-down were defined as markers of hepatocyte clusters.

Taken together these results support specific labelling of renal proximal tubule epithelial cells in the kidney and of hepatocytes in the liver, with enrichment of their transcriptomes in pulled-down RNA and specific depletion for genes associated with other cell types found in these organs. To assess the feasibility of detecting dynamic transcriptional responses using iTag-RNA, we examined whether diet-induced transcriptional reprogramming can be identified in pulled-down poly-A RNA, and to what extent it reflects transcriptional changes observed in whole-tissue input RNA. Differential gene expression analyses using the DEseq package [55] revealed a substantial overlap between diet induced transcriptional reprogramming in the liver as observed in input mRNA to transcriptional reprogramming observed in pull-down libraries (Figure 4g, **Sup. table 5)**. Though the total amount of differentially expressed genes **(DEG)** was roughly 4 fold lower in pull-down vs. input libraries (157 vs. 636 DEG, with an FDR cutoff of less than 0.05 and absolute log2 fold change greater than 1), a 61% overlap (96 DEG) between the two sample sets was detected. This overlap is much higher than expected by chance (chi test <0.0001). Diet induced DEG in both input and pull-down liver RNA demonstrated a significant enrichment for genes regulated by PPARA, LXR and RXR (Figure 4h and **Sup. Table 7)**.

As opposed to liver, diet-induced differential expression in the kidney was limited to 108 transcripts in input poly-A RNA enriched for mitochondrial and ribosomal proteins, whilst pulled-down RNA demonstrated no transcriptional reprograming (Figure 4i-j, **Sup. table 6-7).** These findings may reflect the more complex cellular composition of the kidney and a lack of transcriptional reprogramming in proximal renal epithelial cells.

Taken together, these results provide a proof-of-concept that iTAG-RNA allows isolation of cell-type specific transcriptional responses to environmental challenges. Importantly, with no need for the disruption of the tissue architecture or interference with the cellular microenvironment.

### Hepatocyte derived ccfRNA are detected in Plasma

Given the observed hepatic transcriptional reprogramming following a HFD challenge, we examined whether we can detect labeled hepatocyte derived plasma ccfRNAs, and if the profile of these secreted transcripts changes in response to the dietary challenge. Plasma isolated ccfRNAs are predominantly short / fragmented RNA transcripts with a bi-modal distribution and a major peak smaller than 200bp ([56] **and** sup figure 3g). As already described, with a single dose of HD5EU 2 hours before blood collection we failed to generate libraries from plasma ccfRNA following biotinylation and pull-down. However, with multiple doses of HD5EU administration 6, 4 and 2 hours before blood collection, we were able to generate small RNA libraries following pull-down of plasma ccfRNAs.

Multiple short RNAs were pulled-down in both liver and plasma under HFD and LFD. (Figure 5a, 459/992 for HFD and 234/700 for LFD). This co-occurrence rate is greater than expected by chance for HFD and LFD (Figure 5a, P.V calculated using the SuperExactTest package in R [57], Fold enrichment = HFD: 2.7; LFD = 1.8; P.V. = HFD: 4.3e-104; LFD = 6.42e-21). The identity of pulled-down plasma ccfRNAs varied between dietary challenges (Figure 5b **and Sup. tables 8-10)**, with multiple reads found enriched only under a specific dietary challenge, suggesting that liver secreted ccfRNAs indeed change with dietary interventions.

**Figure 5.**
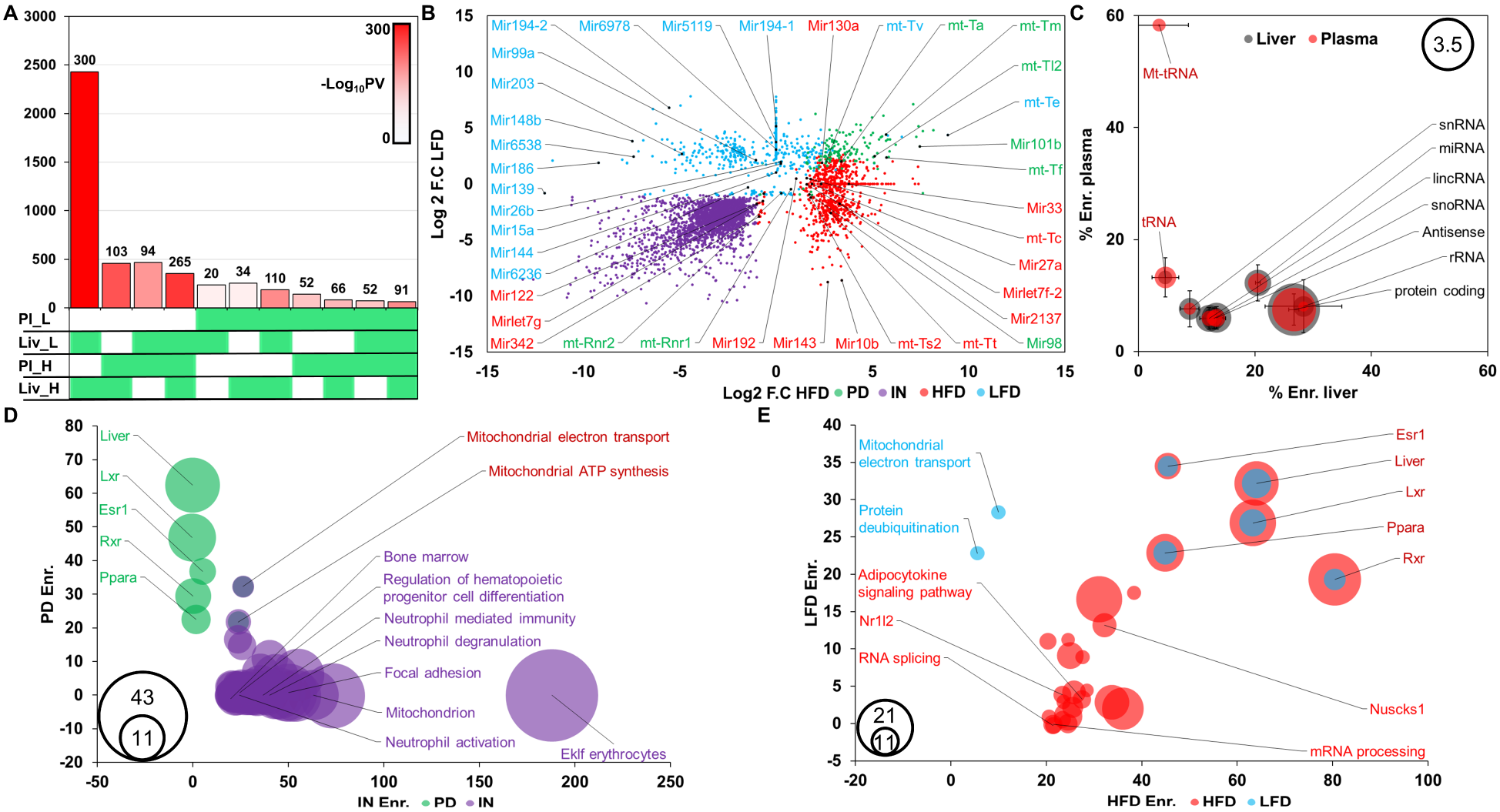
Identification of hepatocyte secreted circulating transcripts in Plasma. **A)** Overlap between pulled-down small RNA transcripts in liver and plasma under HFD and LFD challenge. Number of transcripts per overlap indicated in the y-axis. Color corresponds to –log_10_ of calculated P.V and is annotated in the chart. Gene sets included in each comparison are indicated in green. **B)** Scatter plot annotating selected miRNAs and mt-tRNAs identified as pulled-down in circulating plasma RNA under HFD and LFD conditions. Transcripts are either depleted following pull-down regardless of dietary regime (purple), pulled-down regardless of dietary regime (green) or pulled-down under HFD (red) or LFD (blue) dietary regimes only. **C)** Bubble plot representing the relative percentage of pulled-down transcripts per biotype in liver and plasma libraries, averaged across HFD and LFD regimes. Bubble size represents log10 of actual number of transcripts per bio-type. Error bars for standard deviation between sets. **D)** Bubble plot of GO and gene set enrichment analysis for constitutively pulled-down (green) and depleted (purple) cell-free circulating transcripts. Bubble size proportional to –log_10_ of adj. PV. **E)** Bubble plot of GO and gene set enrichment analysis for cell-free circulating transcripts pulled-down in HFD (red) or LFD (blue). Bubble size proportional to –log_10_ of adj. PV.

Various biotypes are identified in pulled-down liver and plasma short RNA libraries, with the relative proportion of pulled-down transcripts varying between the two. tRNAs, mitochondrial tRNAs and mitochondrial genes are found to be overly represented in pull-down RNA from plasma relative to liver (Figure 5c). This result suggests that the majority of mitochondrial transcripts found in plasma originate predominantly in the liver.

Additional support in favor of the hepatic origin of plasma labeled transcripts can be found in fragments originating from protein coding genes. These protein coding fragments demonstrate a significant enrichment for liver specific and highly expressed genes, whilst transcripts constitutively depleted in pull-down RNA demonstrate an enrichment for bone marrow specific protein coding fragments, genes related to hematopoietic differentiation and genes specific to neutrophil function. (Figure 5d **and sup. table 11)**. This result may suggests that the hematopoietic system is one of the major contributors of circulating RNAs. HFD and LFD specific pulled-down protein coding transcripts demonstrate differential enrichment for annotations including adipocytokine signaling and mitochondrial electron transport respectively (Figure 5e). The significant enrichment found for liver specific protein coding genes among pulled-down transcripts supports the hypothesis that pulled-down ccfRNA transcripts originate in hepatocytes, where Cyp3a4 expression metabolizes HD5EU to 5EU, allowing its incorporation into nascent transcribing RNA.

### Hepatic derived ccfRNA are found in visceral white adipose tissue where they contribute to the small RNA pool and potentially regulate lipid storage

In worms, plants and prokaryotes, extracellular RNA signaling was described to modulate host/pathogen interactions and to orchestrate an adaptive response to environmental stimuli [58-60], and demonstrated the idea that biological systems can exist as holobionts characterized by continuous exchange of genetic (DNA, RNA) material. In mammals, RNA-based intercellular signaling has been described in several settings [4, 9, 12, 61-63], using mostly in-vitro systems or ectopic administration of RNAs to demonstrate RNA transfer and signal transduction. Little is known on the extent of RNA transfer in-vivo and on the role it may play in physiological settings. Two weeks of high-fat diet feeding are sufficient to impair metabolic homeostasis [64], and induce morpho-functional alterations in both liver and visceral adipose tissue [64]. Given the central role liver and adipose tissue play in metabolic control [65], and existing evidence suggesting RNA-based signaling between the tissues [12], we used iTAG-RNA to identify diet-sensitive RNA-based liver-to-adipose signals.

The number of transcripts found to be enriched following pull-down in liver, plasma and VsWAT was 9.7 folds greater than expected by chance for HFD and 6.8 folds greater than expected for LFD (HFD: 264/27, PV=6.34e-180; LFD: 116/17, PV = 3.7e-59 as calculated using the SuperExactTest package in R [57]) **(Figure 6a Sup tables 8-10)**. Focusing on identified hepatic transcribed plasma ccfRNAs, multiple small RNAs could be enriched for in VsWAT. These include miRNAs such as mir-33, mir-10b and mir-130a, in addition to mt-tRNAs (**Figure 6b)**. Among experimentally validated targets of the identified miRNAs (as annotated by miRTarBase [66]), an enrichment is found for proteins associated with the GO term negative regulation of lipid storage (GO:0010888, adj. P.V. = 0.0078, enrichment score = 26.68) such as Abca1, Ppara and Pparg which are regulated by mir-33 [23, 24], mir-10b [25] and mir-130a [26, 67] respectively. Mir-33 is also an identified regulator of the Srebp family of transcription factors that are central in cholesterol and fatty acid synthesis [68, 69] and is in fact encoded within the intron of Srebf2. Total RNA sequencing of VsWAT identified 100 genes to be differentially expressed following acute HFD feeding (28 upregulated / 72 downregulated, **Sup table 12**). As expected given the increased dietary intake of free fatty acids following HFD feeding, downregulation of Srebf1 and of several target genes involved in fatty acid biosynthesis such as FasN, Acaca, and Scd2 is evident (**Figure 6c-d**).

**Figure 6.**
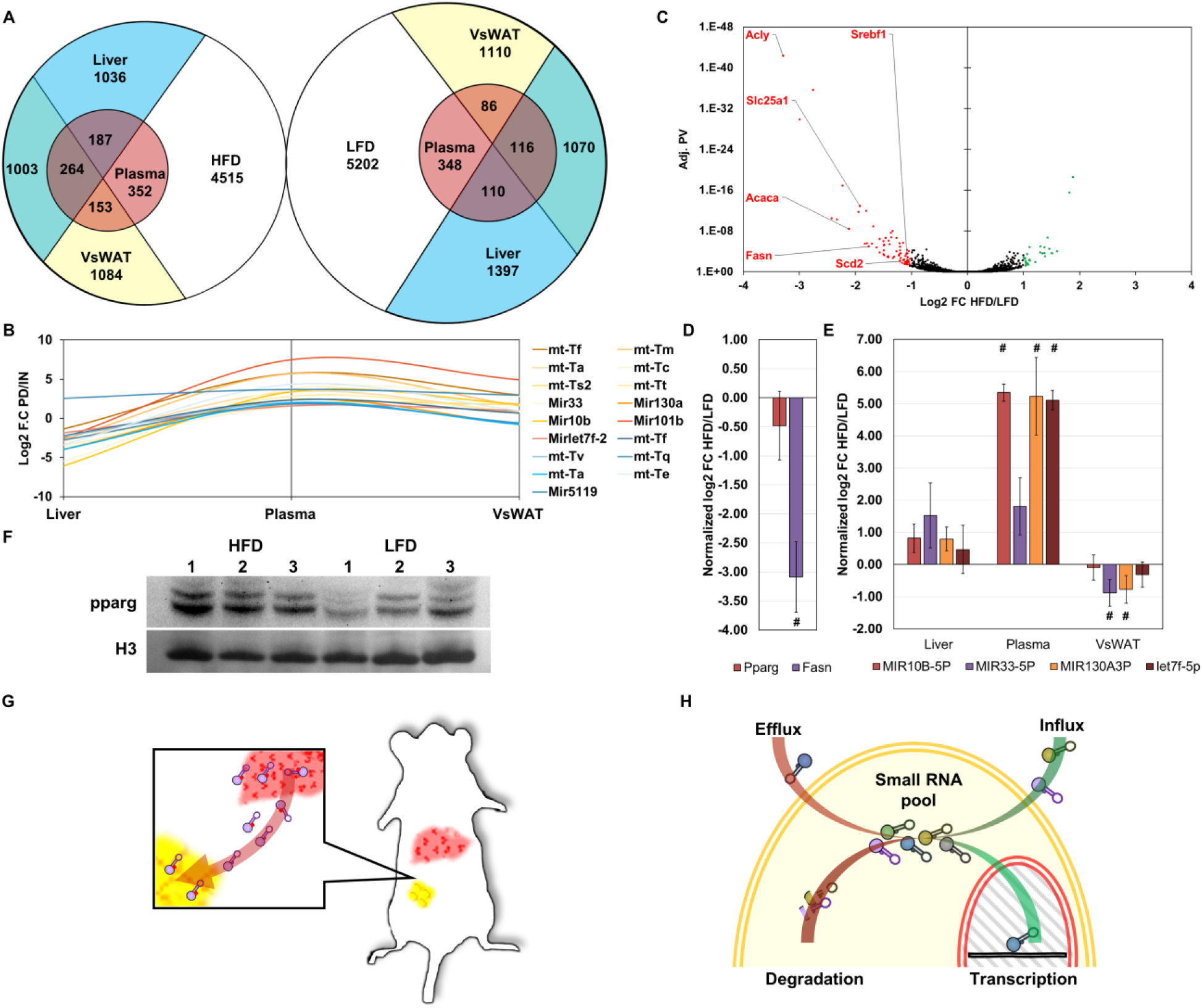
Plasma ccfRNAs can be detected in VsWAT. **A)** Venn diagram demonstrating the overlap between pulled-down transcripts in expression sets from liver plasma and VsWAT for HFD and LFD (Blue = Liver; Red = Plasma; Yellow = VsWAT). **B)** miRNAs and mt-tRNAs that are pulled-down in both plasma and VsWAT (Red = HFD; Blue = LFD). **C)** Volcano plot demonstrating DEG in total RNA sequencing of VsWAT following 2 weeks of HFD or LFD challenge. Red – downregulated in HFD, Green – Upregulated in HFD. **D)** qRT-PCR estimations for differential expression of Pparg and Fasn in VsWAT. # = significant fold change, n=3. **E)** qRT-PCR estimations for differential expression of selected miRNAs. # = significant fold change, n=3. **F)** Western blot for Pparg, modest upregulation of Pparg levels is evident following 2 weeks of HFD. **G)** Depiction of small RNA transfer between liver and adipose tissue. H) A model suggesting that the small RNA pool within a cell results from a balance between transcription and degradation on the one hand and influx and efflux on the other.

## Discussion

In this work we present **iTAG-RNA**, a novel method for targeted in-vivo labelling of global RNA transcription, and use it to identify hepatocyte secreted ccfRNA and their uptake by adipose tissue in-vivo. iTAG-RNA allows labelling of total RNA transcripts in-vivo using two main components: **HD5EU**, a newly designed small molecule that serves as a metabolite for the human CYP3A4 enzyme; and **an existing humanized transgenic mouse model** expressing the human CYP3A4 enzyme under a modified APOE promoter (Figure 1). CYP3A4 catalyzes the oxidative cleavage of an aryl group of the HD5EU molecule, which in turn undergoes spontaneous beta elimination to produce a bio-available 5EU-monophosphate. The existing transgenic mouse model allows in-vivo labelling of hepatocytes and kidney proximal renal epithelial cells (Figure 2), as the tissue expression pattern of the enzyme dictates the site of the small molecule’s metabolism and subsequent RNA labeling. To validate the specificity of HD5EU metabolism, we demonstrate that HD5EU is indeed metabolized in a CYP3A4 dependent manner to 5EU-monophosphate. 5EU is then incorporated in place of uridine into transcribing RNA (Figure 2-3) and allows highly selective RNA precipitation and sequencing of labeled RNAs (Sup. Figure 3 and Figure 4a). Development of new transgenic models would allow for labeling of multiple tissues of choice. Administration of HD5EU allows enrichment for the transcriptional program of proximal renal epithelial cells and hepatocytes in-situ without disruption of the kidney or liver architecture (Figure 4b-g). In addition, environmentally induced transcriptional reprogramming is evident following labelling (Figure 4h-k). As opposed to recently described methods [14, 17, 20, 22] and in keeping with the literature where POLI, POLII and POLIII are demonstrated to incorporate 5EU [27], mRNA and small RNAs of various types including rRNA, tRNA and miRNA are found to be labeled and enriched in pulled-down RNA.

Critically, and uniquely to iTag-RNA, we are also able to enrich for liver derived plasma ccfRNA following an administration of multiple doses of HD5EU (Figure 5). Pulled-down plasma ccfRNAs demonstrate an enrichment for liver derived RNA fragments of protein-coding genes, whilst depleted transcripts demonstrate an enrichment for annotation relating to function and differentiation of the hematopoietic system. Apart from fragments of protein coding genes, liver secreted ccfRNA include various small RNA transcripts such as miRNA, mt-tRNAs and tRNAs. Given the evident enrichment for mitochondrial transcripts following pull-down, our results suggest that hepatocytes are the main source of mitochondrially encoded ccfRNA transcripts in plasma. The functional significance of these tRNA fragments is unclear, but several studies have suggested tRNAs and tRNA fragments can mediate cellular signaling, and that the overall tRNA pool and composition within a cell has functional significance [70].

We continued to explore the possibility that ccfRNAs are taken up by tissues in-vivo to mediate cell-to-cell communication. Focusing on VsWAT we pulled-down and amplified labeled RNA, again following multiple doses of HD5EU **(Figure 6a)**. The transcripts we identified as labeled in VsWAT demonstrated a greater overlap than expected by chance with labeled plasma ccfRNAs or indeed with labeled transcripts in the liver, supporting their hepatic transcriptional origin. These transcripts including miRNAs and mt-tRNAs **(Figure 6b)**, with the identified miRNAs implicated in post-transcriptional regulation of proteins involved in lipid, cholesterol and fatty acid pathways. Together our results directly demonstrate for the first time, in-vivo transfer of a large variety of RNAs including miRNAs between hepatocytes and visceral adipose tissue (**Figure 6g)**, with the identity of transferred RNAs varying following an acute dietary challenge. In light of the observed diversity and scope of RNA transfer between hepatocytes and adipocytes, our results suggest that the pool of small RNAs in a cell in-vivo results from a balance not only between transcription and degradation, but also influx and efflux of small RNAs from the extracellular environment (**Figure 6h)**, with the latter serving as a buffering mechanism for transcriptional and physiological responses in target cells.

To date, the majority of potential ccfRNA biomarkers associated with liver pathologies have been miRNAs [71]. Our findings suggest that fragments of protein coding genes together with mitochondrial tRNAs and mitochondria encoded transcripts can also serve as useful biomarkers for hepatic function. Development of additional genetic models similar to the one used can allow for better transcriptional characterization of distinct cell populations in-vivo, with the added benefit of labelling endogenous ccfRNAs in-vivo. Lastly, HD5EU may prove beneficial in clinical settings. As metabolic activation of HD5EU requires the human Cyp3a4 enzyme, whose expression under physiological conditions is largely limited to the liver (The human protein atlas. 2019. CYP3A4. [ONLINE] Available at: https://www.proteinatlas.org/ENSG00000160868-CYP3A4/tissue/primary+data **[72])**, administration of this small molecule to humans may constitute a novel diagnostic tool allowing assessment of hepatic function by means of a liquid biopsy.

## Methods

### Mice handling

Mice were purchased from Taconic (Taconic USA). All mice were kept in a SPF facility in accordance with the Bavarian Animal law. Mice were fed with a chow / high fat diet / low fat diet as indicated (Rodent Diet with 60 kcal% from fat - Research Diet D12492i, Rodent Diet with 10 kcal% from fat – Research Diet D12450B). 5EU (7848.2, Carl Roth) was solubilized in saline 0.9% NaCl, HD5EU in a 25 % PEG-400, 5% DMSO saline solution. Compounds were administered intraperitoneally at a dose of 0.15 / 0.3_mg/g_ in a total volume of 200µl. First administration was always carried out at ZG-3 to avoid circadian effects. For blood and organ collection, mice were terminally anesthetized with Ketamin/Xylazine at indicated times following drug administration. Heart puncture was performed to collect blood in EDTA coated syringes. Blood was centrifuged at 4.8K rpm for 10’ followed by 12K rpm at 20’ and filtration through a 22uM PES filter (PA59.1, Carl Roth). For isolation of primary hepatocytes, mice were anesthetized and liver perfused through the vena cava with Gibco’s liver perfusion buffer (17701-038, Gibco) and liver digestion buffer (17703-034, Gibco) in accordance with manufacturer’s instructions.

### Cell culture

Isolated primary hepatocytes were counted using the countess automated cell counter (C10227, invitrogen), and plated to a density of 75k/Cm^2^ on Geltrex (A1413201, ThermoFisher) coated coverslips (200 µg/cm^2^) in Williams’ Medium E (A12176, Gibco) supplemented with Gibco’s Primary Hepatocyte Maintenance Supplements (CM4000, Gibco). 24 hours following plating, cells were treated with 1mM of 5EU or HD5EU for 8 hours. Azamulin (SML0485, Sigma Aldrich) was added at a concentration of 20µM for 30 minutes before addition of indicated compounds to a final concentration of 10µM for the length of the treatment.

### Tissue processing and imaging

Tissues were fixed in a neutral buffered 10% formalin solution (HT501128, Sigma) for 48 hours before dehydration and embedding in paraffin in accordance with published protocols. 4µm sections were cut on a Leica microtome (RM2165, Institute of Experimental Genetics), rehydrated and stained using the Click-iT™ RNA Alexa Fluor™ 594 Imaging Kit (C10330, ThermoFisher) in accordance with manufacturer’s instructions. Mounting was done with Vectashield hardset antifade mounting medium with DAPI (H-1500, Vector Laboratories). Imagining was done using a Laser Scanning Confocal Microscope (Olympus Fluoview 1200, Institute for Diabetes and Cancer, Neuherberg, Germany) equipped with an Olympus UPlanSApo 60x 1.35 Oil immersion objective.

### Western Blot

Tissues were homogenized using a Miltenyi gentleMACS Dissociator (Miltenyi biotec) in RIPA buffer supplemented with protease (S8820, Sigma Aldrich) and phosphatase (88667, Thermo Fisher) inhibitors. Protein concentration was measured using a standard Bradford assay reagent (B6916, Sigma Aldrich).40µg total protein were loaded per samples on a pre-cast gradient 4-12% gel (NW04120, Invitrogen). Proteins were transferred to a PVDF membrane (ISEQ00010, Merck Millipore) blocked and blotted using the iBind system (SLF1020, Invitrogen) with primary anti-CYP3A4 (MA5-17064, Thermo Fisher), anti-Phospho-p53 S392 (#9281, cell signaling), anti-total-p53 (#2524, cell signaling), anti-cleaved Caspase-3 (ab214430, abcam), anti-total Caspase-3 (#ab184787, abcam), anti-pparg (MA5-14889, Thermo Fisher), anti-histone H3 (4499s, cell signalling) and anti-HSP90 (SC-7949, Santa-Cruz), and secondary IgG HRP (7076 and 7074, Cell Signaling)

### RNA extraction, qRT-PCR, pull-down and library construction

Plasma RNA was extracted using TRI Reagent BD (T3809, Sigma Aldrich), in accordance with manufacturer’s instructions. RNA from tissues was extracted using NucleoZOL (740404.200, Macherey-Nagel) reagent, in accordance with manufacturer’s instructions. For qRT-PCR, reverse transcription was conducted using the high-Capacity cDNA Reverse Transcription Kit (4368814, Applied Biosystems), in accordance with the manufacturer’s instructions. Real-time was carried out on a quant-studio 6 flex (applied biosystems) with SYBR Green PCR Master Mix (#4309155 applied biosystems) and primers; hCyp3-F: TTGGCATGAGGTTTGCTCTC; hCyp3-R: ACAACGGGTTTTTCTGGTTG; Pparg-F: AGATTCTCCTGTTGACCCAGAG; Pparg-R: AGCTGATTCCGAAGTTGGTG; Fasn-F: CTGCTGTTGGAAGTCAGCTATG; Fasn-R: ATGCCTCTGAACCACTCACAC; Actin-F: CACAGCTTCTTTGCAGCTCCT; Actin-R: CAGCAGTGCAATGTTAAAAGG; qRT-PCR for miRNAs was conducted as previously described [73] and primers designed with miRprimer2 [74]. miR-7f-5p-F: CGCAGTGAGGTAGTAGATTG; miR-7f-5p-R: CAGGTCCAGTTTTTTTTTTTTTTTAAC; miR-130a-3p-F: CAGCAGTGCAATGTTAAAAGG; miR-130a-3p-R: CAGGTCCAGTTTTTTTTTTTTTTTATG; miR-33-5p-F: CGCAGGTGCATTGTAGT; miR-33-5p-F:GTCCAGTTTTTTTTTTTTTTTGCAAT; miR-10b-5p-F CAGTACCCTGTAGAACCGA; miR-10b-5p-R:GGTCCAGTTTTTTTTTTTTTTTCAG; Pull-down of 5EU labeled RNA was done using the Click-it Nascent RNA Capture Kit (C10365, Thermo Fisher). 10µg total / small RNA was used as input from tissues, 200ng plasma RNA was used as input for RNA pull-down from blood. RNA was used as template for library construction using the CATS mRNA/small RNA kit (C05010043 and C05010040, Diagenode) with slight modifications to protocol i.e.; Poly-A selection and RNA fragmentation were performed before biotinylation and pull-down of 5EU. 14 cycles of amplification for RNA from tissues, 20 for RNA from blood. 10ng of RNA was used for Input. For MS analysis, 10µg long / short RNA was used. RNA was digested as described in Sue et. al. [41].

### UPLC-UHR-ToF-MS analysis

Mass spectrometric analysis of nucleotides from RNA was performed on a Waters Acquity UPLC (Waters, Eschborn, Germany) coupled to a Bruker maXis UHR-ToF-MS (Bruker Daltonic, Bremen, Germany). Separation was performed on a Thermo Hypersil Gold column (150 x 1.0 mm, 3 µm, 25003-151030, Thermo Fisher) using a multistep gradient with 100% water and 100% ACN, both with 0.1% formic acid. Gradient conditions were as followed: 0-6 min 0% B, 6-7.65 min linear increase to 1% B, 7.65 to 10 min linear increase to 6% B, 10 to 12 min linear increase to 50% B, 12 to 14 min linear increase to 75% B, 14 to 17 min isocratic hold of 75% B, 17 to 17.5 min return to initial conditions. Column temperature was 36°C and flow rate was set to 0.09 ml/min. Before each run the column was re-equilibrated for 3 minutes with starting conditions. High mass accuracy was achieved by infusion of 1:4 diluted ESI low concentration tune mix (Agilent Technologies, Waldbronn, Germany) at the start of each chromatographic run. Each analysis was internally recalibrated using the tune mix peak at the beginning of the chromatogram using a custom VB script within Bruker DataAnalysis 4.0 (Bruker Daltonic, Bremen, Germany). Quantitative analysis was performed in Bruker QuantAnalysis 4.0 (Bruker Daltonic, Bremen, Germany). High Resolution-Extracted Ion Chromatograms (HR-EICs) were created around each precursor mass +/- 0.005 Da. Chromatograms were smoothed and peak areas were used for quantification. In case of 5EU additional quantification was performed on a validated in-source fragment [M-132+H]^+^.

### Bioinformatic analysis

RNA Libraries were sequenced on an Illumina HiSeq 2500 instrument (**IGA Technology Services Srl, Italy)** at 75bp single-ended. Adaptors were trimmed in accordance with the CATS sequencing kit manual. Reads were aligned to the mouse mm10 genome using the STAR aligner [43] and a reference transcript gtf file from ensamble modified to contain tRNA transcripts as annotated by GtRNAdb [75].

For the detection of differentially regulated genes between HFD and LFD the Deseq2 package was used [55]. Pull-down enrichment analysis was conducted using the NOISeq package. Transcripts with expression values smaller than a cpm of 1 and a coefficient of variation greater than 300 were filtered out prior to tmm normalization and enrichment analysis. GO and gene set enrichment was calculated using the Enrichr tool [45].

## Acknowledgments

We would like to thank Dr. Julia Calzada-Wack, Jacqueline Mueller and Marion Fisch for their kind assistance with tissue processing and sectioning. We thank Dr. Anja Zeigerer for access to and assistance with the confocal microscope. This work has been supported by the German Diabetes Research Center (DZD NEXT Grant 2019) to T.R., and by the ERC Recognition Award from the Helmholtz Research Center Munich to T.R. The authors thank the Helmholtz Association and the German Diabetes Research Center for funding the positions of G.R., D.J., L.M., S.F., T.A. and T.R.

## Author contributions

Conceptualization D.J.; Methodology D.J. and W.M.; Investigation D.J., L.M., G.R., A.T., S.F. and W.M.; Writing – Original Draft D.J., W.M., and T.R.; Writing – Review & Editing D.J., H.A.M, W.M. and T.R.; Funding Acquisition H.A.M and T.R.; Resources; H.A.M, W.M. and T.R.; Supervision T.R.

## Supplementary figure legends

**Supplementary Figure 1.**
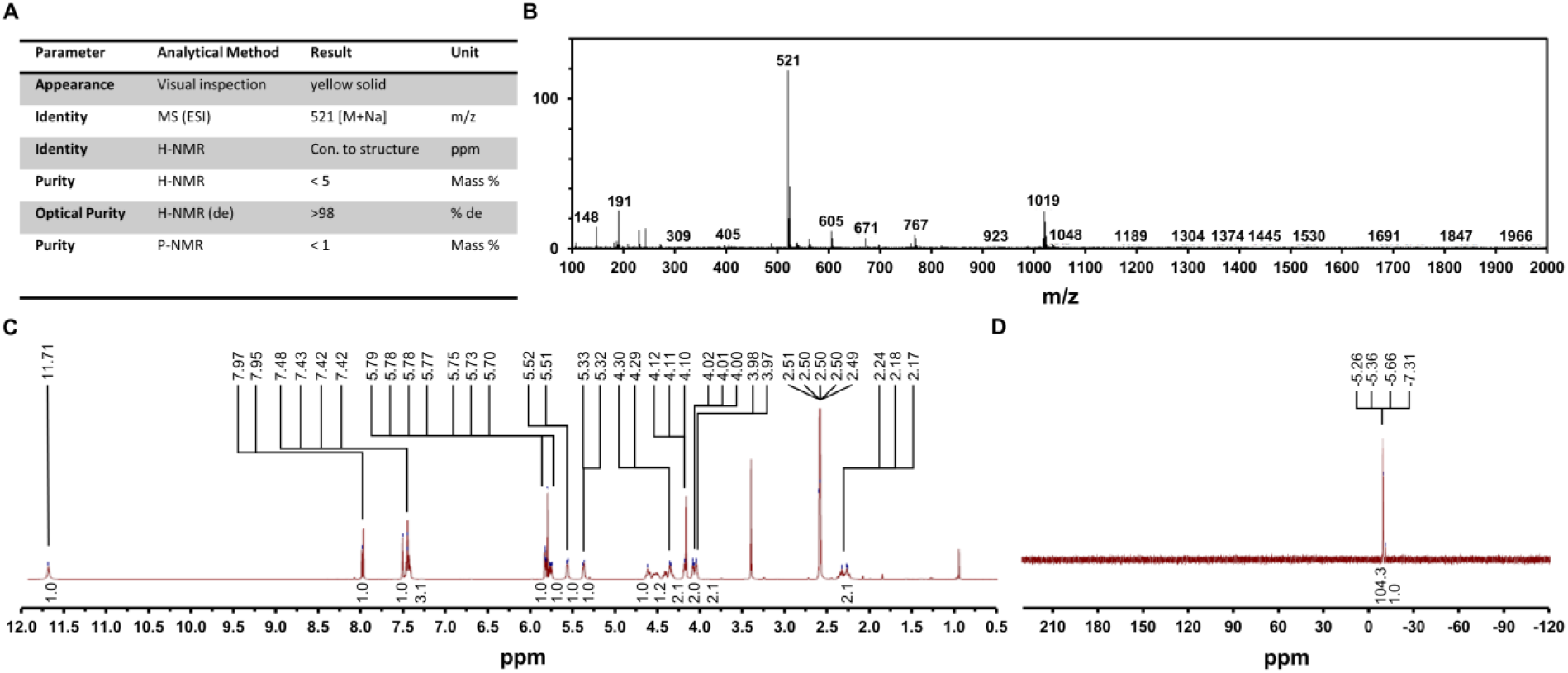
Quality assessment of HD5EU synthesis. **a)** Summary table listing all examined parameters and methodologies used to validate the structural conformity of the molecule and the purity of the product. **b)** MS Base Peak at 520.95 corresponding to M+NA^+^ and at 1019 for 2M+NA^+^. **c)** ^1^H-NMR conformity to structure. 513 MHz. DMSO as solvent. **d)** ^31^p-NMR measurement for purity. 202.46 MHz. DMSO as solvent.

**Supplementary Figure 2.**
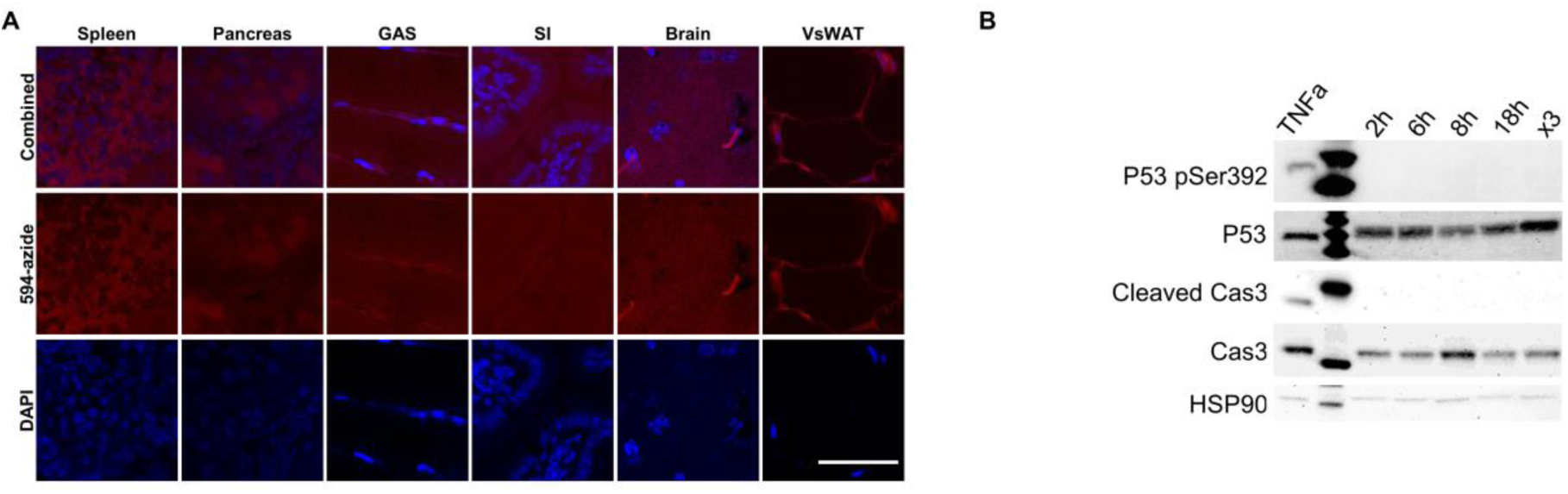
Negative controls for in-vivo tissue staining. **a)** Click-it staining in tissues collected from saline treated animals. Scale Bar = 50µM. **b)** W.B. validating HD5EU safety. Lack of p53 activation and downstream Caspase cleavage at multiple time points following administrations of HD5EU and following consecutive administration of HD5EU. MEFs treated with TNFa for 16 hours serve as positive controls for the western blot staining.

**Supplementary Figure 3.**
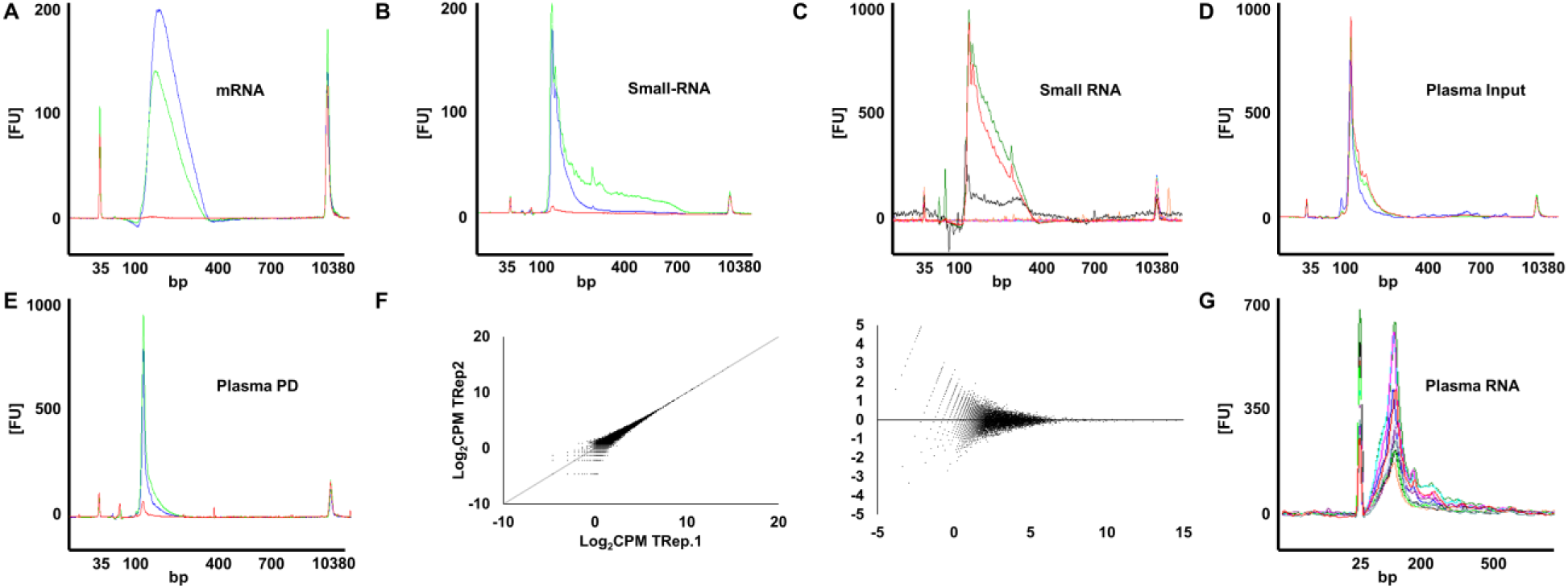
Specificity and reproducibility of RNA pull-down, library construction and sequencing. **A)** Bio-analyzer plot for mRNA Pull-down libraries. Blue - HD5EU labelled liver, Green - HD5EU labelled kidney, Red - saline treated liver. We consistently failed to generated libraries from non-labeled RNA subject to biotinylation and pull-down. **B)** Bio-analyzer plot for small RNA pull-down libraries. Blue – HD5EU labelled liver, Green - HD5EU labelled kidney, Red – Saline treated liver. **C)** Consistent failures in library generation from small RNA (<200bp) pull-down in multiple tissue following 2h HD5EU treatment (Testis = orange, VsWAT = pink and blue = Spleen). Input RNA generates expected library amplicons (Black, red and green). **D)** Bio-analyzer plots for small RNA input libraries from plasma. **E)** Bio-analyzer plots for small RNA pull-down libraries from plasma following multiple injections. Unlabeled RNA in Red is amplified as input but fails to amplify following Pull-down. **F)** Scatter plot and MA plot for two technical replicates from HD5EU labeled hepatocyte samples. Pearson correlation coefficient = 0.95. **G)** Plasma ccfRNAs’ size distribution.

## Supplementary Tables

**Supplementary table 1.** Identification and separation of various modified and unmodified transcripts in LC-MS.

**Supplementary table 2.** Noiseq-bio analysis for the detection of enriched and depleted transcripts following pull-down of labelled poly-A RNA in liver and kidney.

**Supplementary table 3.** GO and gene set enrichment analysis for kidney pulled-down and depleted poly-A RNA.

**Supplementary table 4.** GO and gene set enrichment analysis for liver pulled-down and depleted poly-A RNA.

**Supplementary table 5.** Differential expression analysis using Deseq2, for the detection of dietary induced DEG in liver, input and pull-down poly-A RNA libraries.

**Supplementary table 6.** Differential expression analysis using Deseq2, for the detection of dietary induced DEG in kidney, input and pull-down poly-A RNA libraries.

**Supplementary table 7.** GO and gene set enrichment analysis on dietary induced differentially expressed genes in liver and kidney pull-down or depleted poly-A RNA.

**Supplementary table 8.** Noiseqbio differential pull-down analysis for liver, plasma and VsWAT under HFD regime. TMM normalized mean expression values.

**Supplementary table 9.** Noiseqbio differential pull-down analysis for liver, plasma and VsWAT under LFD regime. TMM normalized mean expression values.

**Supplementary table 10.** SuperExacttest package output. Overlaps between different sets and calculated P.V.

**Supplementary table 11** GO and gene set enrichment analysis on plasma pulled-down protein coding genes.

**Supplementary table 12** Differential expression analysis using Deseq2 package, for the detection of dietary induced DEG in total VsWAT RNA.

## Reference

1. Murillo, O.D., et al., exRNA Atlas Analysis Reveals Distinct Extracellular RNA Cargo Types and Their Carriers Present across Human Biofluids. Cell, 2019. 177(2): p. 463–477 e15.

2. Yeri, A., et al., Total Extracellular Small RNA Profiles from Plasma, Saliva, and Urine of Healthy Subjects. Sci Rep, 2017. 7: p. 44061.

3. Dror, S., et al., Melanoma miRNA trafficking controls tumour primary niche formation. Nat Cell Biol, 2016. 18(9): p. 1006–17.

4. Castano, C., et al., Obesity-associated exosomal miRNAs modulate glucose and lipid metabolism in mice. Proc Natl Acad Sci U S A, 2018. 115(48): p. 12158–12163.

5. Wortzel, I., et al., Exosome-Mediated Metastasis: Communication from a Distance. Dev Cell, 2019. 49(3): p. 347–360.

6. Schwarzenbach, H., et al., Clinical relevance of circulating cell-free microRNAs in cancer. Nature Reviews Clinical Oncology, 2014. 11: p. 145.

7. Gilad, S., et al., Serum microRNAs are promising novel biomarkers. PLoS One, 2008. 3(9): p. e3148.

8. Pegtel, D.M., et al., Functional delivery of viral miRNAs via exosomes. Proc Natl Acad Sci U S A, 2010. 107(14): p. 6328–33.

9. Vickers, K.C., et al., MicroRNAs are transported in plasma and delivered to recipient cells by high-density lipoproteins. Nat Cell Biol, 2011. 13(4): p. 423–33.

10. Pastuzyn, E.D., et al., The Neuronal Gene Arc Encodes a Repurposed Retrotransposon Gag Protein that Mediates Intercellular RNA Transfer. Cell, 2018. 172(1-2): p. 275–288 e18.

11. Kosaka, N., et al., Secretory mechanisms and intercellular transfer of microRNAs in living cells. J Biol Chem, 2010. 285(23): p. 17442–52.

12. Thomou, T., et al., Adipose-derived circulating miRNAs regulate gene expression in other tissues. Nature, 2017. 542(7642): p. 450–455.

13. Sharma, U., et al., Small RNAs Are Trafficked from the Epididymis to Developing Mammalian Sperm. Dev Cell, 2018. 46(4): p. 481–494 e6.

14. Gay, L., et al., Mouse TU tagging: a chemical/genetic intersectional method for purifying cell type-specific nascent RNA. Genes Dev, 2013. 27(1): p. 98–115.

15. Herzog, V.A., et al., Thiol-linked alkylation of RNA to assess expression dynamics. Nat Methods, 2017. 14(12): p. 1198–1204.

16. Chatzi, C., et al., Transcriptional Profiling of Newly Generated Dentate Granule Cells Using TU Tagging Reveals Pattern Shifts in Gene Expression during Circuit Integration. eNeuro, 2016. 3(1).

17. Miller, M.R., et al., TU-tagging: cell type-specific RNA isolation from intact complex tissues. Nat Methods, 2009. 6(6): p. 439–41.

18. Ghosh, A.C., et al., UPRT, a suicide-gene therapy candidate in higher eukaryotes, is required for Drosophila larval growth and normal adult lifespan. Sci Rep, 2015. 5: p. 13176.

19. Maquat, L.E. and M. Kiledjian, RNA turnover in eukaryotes: nucleases, pathways and analysis of mRNA decay. Preface. Methods Enzymol, 2008. 448: p. xxi–xxii.

20. Hida, N., et al., EC-tagging allows cell type-specific RNA analysis. Nucleic Acids Res, 2017. 45(15): p. e138.

21. Song, A.J. and R.D. Palmiter, Detecting and Avoiding Problems When Using the Cre-lox System. Trends Genet, 2018. 34(5): p. 333–340.

22. Alberti, C., et al., Cell-type specific sequencing of microRNAs from complex animal tissues. Nat Methods, 2018. 15(4): p. 283–289.

23. Marquart, T.J., et al., miR-33 links SREBP-2 induction to repression of sterol transporters. Proc Natl Acad Sci U S A, 2010. 107(27): p. 12228–32.

24. Rayner, K.J., et al., MiR-33 contributes to the regulation of cholesterol homeostasis. Science, 2010. 328(5985): p. 1570–3.

25. Zheng, L., et al., Effect of miRNA-10b in regulating cellular steatosis level by targeting PPAR-alpha expression, a novel mechanism for the pathogenesis of NAFLD. J Gastroenterol Hepatol, 2010. 25(1): p. 156–63.

26. Lee, E.K., et al., miR-130 suppresses adipogenesis by inhibiting peroxisome proliferator-activated receptor gamma expression. Mol Cell Biol, 2011. 31(4): p. 626–38.

27. Jao, C.Y. and A. Salic, Exploring RNA transcription and turnover in vivo by using click chemistry. Proc Natl Acad Sci U S A, 2008. 105(41): p. 15779–84.

28. Best, M.D., Click chemistry and bioorthogonal reactions: unprecedented selectivity in the labeling of biological molecules. Biochemistry, 2009. 48(28): p. 6571–84.

29. Hagemeijer, M.C., et al., Visualizing coronavirus RNA synthesis in time by using click chemistry. J Virol, 2012. 86(10): p. 5808–16.

30. Gierlich, J., et al., Click chemistry as a reliable method for the high-density postsynthetic functionalization of alkyne-modified DNA. Org Lett, 2006. 8(17): p. 3639–42.

31. Meyer, J.P., et al., Click Chemistry and Radiochemistry: The First 10 Years. Bioconjug Chem, 2016. 27(12): p. 2791–2807.

32. Pradere, U., et al., Synthesis of nucleoside phosphate and phosphonate prodrugs. Chem Rev, 2014. 114(18): p. 9154–218.

33. Erion, M.D., et al., Design, synthesis, and characterization of a series of cytochrome P(450) 3A-activated prodrugs (HepDirect prodrugs) useful for targeting phosph(on)ate-based drugs to the liver. J Am Chem Soc, 2004. 126(16): p. 5154–63.

34. Boyer, S.H., et al., Synthesis and characterization of a novel liver-targeted prodrug of cytosine-1-beta-D-arabinofuranoside monophosphate for the treatment of hepatocellular carcinoma. J Med Chem, 2006. 49(26): p. 7711–20.

35. Erion, M.D., et al., Targeting thyroid hormone receptor-beta agonists to the liver reduces cholesterol and triglycerides and improves the therapeutic index. Proc Natl Acad Sci U S A, 2007. 104(39): p. 15490–5.

36. Reddy, K.R., et al., Pradefovir: a prodrug that targets adefovir to the liver for the treatment of hepatitis B. J Med Chem, 2008. 51(3): p. 666–76.

37. van Herwaarden, A.E., et al., Midazolam and cyclosporin a metabolism in transgenic mice with liver-specific expression of human CYP3A4. Drug Metab Dispos, 2005. 33(7): p. 892–5.

38. van Herwaarden, A.E., et al., Knockout of cytochrome P450 3A yields new mouse models for understanding xenobiotic metabolism. J Clin Invest, 2007. 117(11): p. 3583–92.

39. Simonet, W.S., et al., A far-downstream hepatocyte-specific control region directs expression of the linked human apolipoprotein E and C-I genes in transgenic mice. J Biol Chem, 1993. 268(11): p. 8221–9.

40. Stresser, D.M., et al., Highly selective inhibition of human CYP3Aa in vitro by azamulin and evidence that inhibition is irreversible. Drug Metab Dispos, 2004. 32(1): p. 105–12.

41. Su, D., et al., Quantitative analysis of ribonucleoside modifications in tRNA by HPLC-coupled mass spectrometry. Nat Protoc, 2014. 9(4): p. 828–41.

42. Williams, L.M., et al., The development of diet-induced obesity and glucose intolerance in C57BL/6 mice on a high-fat diet consists of distinct phases. PLoS One, 2014. 9(8): p. e106159.

43. Dobin, A., et al., STAR: ultrafast universal RNA-seq aligner. Bioinformatics, 2013. 29(1): p. 15–21.

44. Tarazona, S., et al., Data quality aware analysis of differential expression in RNA-seq with NOISeq R/Bioc package. Nucleic Acids Res, 2015. 43(21): p. e140.

45. Chen, E.Y., et al., Enrichr: interactive and collaborative HTML5 gene list enrichment analysis tool. BMC Bioinformatics, 2013. 14: p. 128.

46. Coudrier, E., D. Kerjaschki, and D. Louvard, Cytoskeleton organization and submembranous interactions in intestinal and renal brush borders. Kidney Int, 1988. 34(3): p. 309–20.

47. Lin, G., E. Finger, and J.C. Gutierrez-Ramos, Expression of CD34 in endothelial cells, hematopoietic progenitors and nervous cells in fetal and adult mouse tissues. Eur J Immunol, 1995. 25(6): p. 1508–16.

48. Fina, L., et al., Expression of the CD34 gene in vascular endothelial cells. Blood, 1990. 75(12): p. 2417–26.

49. Liu, Y., D.K. Qiu, and X. Ma, Liver X receptors bridge hepatic lipid metabolism and inflammation. J Dig Dis, 2012. 13(2): p. 69–74.

50. Kersten, S., et al., Peroxisome proliferator-activated receptor alpha mediates the adaptive response to fasting. J Clin Invest, 1999. 103(11): p. 1489–98.

51. Tyagi, S., et al., The peroxisome proliferator-activated receptor: A family of nuclear receptors role in various diseases. J Adv Pharm Technol Res, 2011. 2(4): p. 236–40.

52. Everett, L., A. Galli, and D. Crabb, The role of hepatic peroxisome proliferator-activated receptors (PPARs) in health and disease. Liver, 2000. 20(3): p. 191–199.

53. Park, J., et al., Single-cell transcriptomics of the mouse kidney reveals potential cellular targets of kidney disease. Science, 2018. 360(6390): p. 758–763.

54. MacParland, S.A., et al., Single cell RNA sequencing of human liver reveals distinct intrahepatic macrophage populations. Nat Commun, 2018. 9(1): p. 4383.

55. Love, M.I., W. Huber, and S. Anders, Moderated estimation of fold change and dispersion for RNA-seq data with DESeq2. Genome Biol, 2014. 15(12): p. 550.

56. Srinivasan, S., et al., Small RNA Sequencing across Diverse Biofluids Identifies Optimal Methods for exRNA Isolation. Cell, 2019. 177(2): p. 446–462 e16.

57. Wang, M., Y. Zhao, and B. Zhang, Efficient Test and Visualization of Multi-Set Intersections. Sci Rep, 2015. 5: p. 16923.

58. Liu, H., et al., Escherichia coli noncoding RNAs can affect gene expression and physiology of Caenorhabditis elegans. Nat Commun, 2012. 3: p. 1073.

59. Weiberg, A., et al., Fungal small RNAs suppress plant immunity by hijacking host RNA interference pathways. Science, 2013. 342(6154): p. 118–23.

60. Timmons, L. and A. Fire, Specific interference by ingested dsRNA. Nature, 1998. 395(6705): p. 854.

61. Rivkin, M., et al., Inflammation-Induced Expression and Secretion of MicroRNA 122 Leads to Reduced Blood Levels of Kidney-Derived Erythropoietin and Anemia. Gastroenterology, 2016. 151(5): p. 999–1010 e3.

62. Chai, C., et al., Metabolic Circuit Involving Free Fatty Acids, microRNA 122, and Triglyceride Synthesis in Liver and Muscle Tissues. Gastroenterology, 2017. 153(5): p. 1404–1415.

63. Rechavi, O., et al., Cell contact-dependent acquisition of cellular and viral nonautonomously encoded small RNAs. Genes Dev, 2009. 23(16): p. 1971–9.

64. Lee, Y.S., et al., Inflammation is necessary for long-term but not short-term high-fat diet-induced insulin resistance. Diabetes, 2011. 60(10): p. 2474–83.

65. Hotamisligil, G.S., Inflammation and metabolic disorders. Nature, 2006. 444(7121): p. 860–7.

66. Chou, C.H., et al., miRTarBase update 2018: a resource for experimentally validated microRNA-target interactions. Nucleic Acids Res, 2018. 46(D1): p. D296–D302.

67. Portius, D., C. Sobolewski, and M. Foti, MicroRNAs-Dependent Regulation of PPARs in Metabolic Diseases and Cancers. PPAR Res, 2017. 2017: p. 7058424.

68. Horie, T., et al., MicroRNA-33 regulates sterol regulatory element-binding protein 1 expression in mice. Nat Commun, 2013. 4: p. 2883.

69. Shimano, H., SREBPs: physiology and pathophysiology of the SREBP family. FEBS J, 2009. 276(3): p. 616–21.

70. Kirchner, S. and Z. Ignatova, Emerging roles of tRNA in adaptive translation, signalling dynamics and disease. Nat Rev Genet, 2015. 16(2): p. 98–112.

71. Enache, L.S., et al., Circulating RNA molecules as biomarkers in liver disease. Int J Mol Sci, 2014. 15(10): p. 17644–66.

72. Thul, P.J., et al., A subcellular map of the human proteome. Science, 2017. 356(6340).

73. Balcells, I., S. Cirera, and P.K. Busk, Specific and sensitive quantitative RT-PCR of miRNAs with DNA primers. BMC Biotechnol, 2011. 11: p. 70.

74. Busk, P.K., A tool for design of primers for microRNA-specific quantitative RT-qPCR. BMC Bioinformatics, 2014. 15: p. 29.

75. Chan, P.P. and T.M. Lowe, GtRNAdb 2.0: an expanded database of transfer RNA genes identified in complete and draft genomes. Nucleic Acids Res, 2016. 44(D1): p. D184–9.

